# Deciphering efficient metabolic pathways in an anaerobic gut fungus coupled with a syntrophic methanogen

**DOI:** 10.64898/2026.10.04.756520

**Authors:** Akshay Joshi, Fabian Meyer, Lona Mosberger, Katharina Cécile Schmid Luedi, Heribert Insam, Gareth Wyn Griffith, Sabine Marie Podmirseg, Rolf Warthmann

## Abstract

Anaerobic gut fungi (AGF) are an integral part of the herbivores’ gut microbiome, primary decomposers of lignocellulose, and play a crucial role in establishing foundational links of syntrophic associations. The syntrophy with methanogens appears beneficial for mitigating feedback inhibition and results in the augmentation of AGF activity. However, the thermodynamic advantages of the presence of methanogens to AGF remain poorly understood. This study explores the preferred pathways adopted by *Cyllamyces* sp. and *Methanobrevibacter olleyae*, using ^13^C-glucose as a substrate and spiked with malate and pyruvate in different experimental set-ups. The results indicated reduced biomass formation by *Cyllamyces* sp. in monoxenic culture with *Methanobrevibacter olleyae* compared to axenic AGF culture supplemented with glucose. Additionally, metabolic analyses revealed the pyruvate:ferredoxin oxidoreductase (PFO) as a key contributor to pyruvate metabolism. Malate supplementation notably enhanced the growth of *Cyllamyces* sp. and *M. olleyae*, whereas pyruvate supplementation enhanced the growth of *Cyllamyces* sp. only in axenic culture. This investigation provides insights into the preferred metabolic pathways operating in the hydrogenosomes of axenic anaerobic gut fungi (AGF) cultures and monoxenic AGF-methanogen co-cultures.

## 1. Introduction

Syntrophic interactions are widespread in anaerobic microbial ecosystems and involve metabolically interdependent microorganisms that cooperatively degrade complex matter through the exchange of metabolic intermediates. These interactions often result in mutual benefits for the partners involved, enabling energy conservation, and driving efficient degradation and enhanced energy production. This intricate process contributes to ecosystem balance, fostering harmonious coexistence among diverse microbial guilds in challenging environments [1].

Anaerobic gut fungi (AGF), belonging to the phylum *Neocallimastigomycota*, are prominent members of the gut microbiota of herbivores [2–4]. They are also known as early-stage degraders of lignocellulosic biomass ingested by the host animal, as they contribute significantly to the degradation process through enzymatic hydrolysis and physical disruption. The metabolites produced by AGF benefit syntrophic microbes with energy and nutrients.

Over the past decade, AGF have attracted increasing research interest due to their exceptional ability to degrade lignocellulosic biomass through the production of extracellular (hemi)cellulolytic enzymes. Their potential involvement in enhancing biogas production alongside syntrophic associations has also drawn significant interest [5–7].

Methanogens and anaerobic bacteria are known to synergistically cooperate with AGF and to play an important role in interspecies electron transfer, preventing end-product inhibition in anaerobic gut fungi [8,9]. The lignocellulose degrading activity, growth, and sugar utilization rate of AGF were observed to be increased in the presence of methanogens [10–12].

Methanogenic archaea can utilize a diverse range of substrates and thus classify mainly into three groups: hydrogenotrophic-, acetoclastic-, and methylotrophic-methanogens. Notably, hydrogenotrophic methanogens often constitute two-thirds of the total methanogenic population in various anoxic environments [13,14]. The hydrogenotrophic methanogens play a crucial role in scavenging hydrogen, produced by biomass-hydrolyzing microorganisms through interspecies hydrogen transfer. These methanogens thrive in the rumen, where volatile fatty acids are rapidly absorbed via the rumen epithelium to provide energy for the animal, thus making them less available for acetoclastic or methylotrophic methanogens.

Most methanogens hitherto cultured from the rumen belong to the families *Methanobacteriaceae*–comprising *Methanobacterium, Methanosphaera*, and *Methanobrevibacter* (sensu stricto, e.g. *M. ruminantium*), and newly delineated genus *Methanocatella* (e.g. *M. gottschalkii*), *Methanoculleus* (*Methanomicrobiaceaea*), and *Methanosarcina* (*Methanosarcinaceae*) [15,16]. In the majority of cases, *Methanobrevibacter* forms the primary syntrophic association with AGF, scavenging H_2_, CO_2,_ and formate [15]. Besides its abundance in the rumen’s archaeal community, the reason for the consistent presence of *Methanobrevibacter* spp. in syntrophic association with AGF merits further investigation. The gut of herbivores reveals another syntrophic interaction of rumen ciliates and methanogens, forming a distinctive endosymbiotic association. Both rumen ciliates and AGF share similarities in producing ATP through substrate-level phosphorylation located in hydrogenosomes, resulting in the formation of similar end products [17][18]. Despite their similarity and syntrophic association with archaea, reports on the specific type of syntrophy between AGF and methanogens remain insufficient.

Anaerobic gut fungi exhibit a complex mixed acid fermentation in the cytosol which leads to energy production both in the cytosol and the hydrogenosome through substrate-level phosphorylation. The hydrogenosome plays a pivotal role in AGF metabolism, serving as an ATP-producing energy unit. The presence of methanogens enhances the lignocellulolytic activity of AGF and alters the metabolic shift from cytosolic acid production towards more intra-hydrogenosomic H_2_, CO_2_, and formate production [10–12].

Syntrophic association with methanogens can influence AGF metabolism and enhance electron flow via hydrogenosomes, thus mitigating feedback inhibition, promoting substrate utilization, increasing ATP production and ultimately boosting AGF activity. However, the hydrogenosomal pathway is also known to shift the cytosolic metabolism of malate and pyruvate, depending on the substrate being utilized for energy production [19]. The energy and end-products driven from these pathways are different and could influence the growth of methanogens. Specifically, pyruvate metabolism only produces formate and acetate, whereas malate metabolism involves CO_2_ in exchange, with the production of H_2_ and formate, and acetate, makes it more favorable for methanogens. Future work should resolve which metabolic pathway is favored in hydrogenosomes in the presence of methanogens would provide insight into the thermodynamic benefits to AGF of such syntrophic interactions.

To elucidate this mechanism, we conducted a comparative growth analysis of *Cyllamyces* sp. RWF43N3 in axenic culture and monoxenic culture with the naturally occurring syntroph *Methanobrevibacter olleyae* RWM43N3. To trace the flow of carbon through intermediate and end-products of AGF metabolism, cultures were grown on a labeled ^13^C_6_-glucose. To further resolve which pathway was favored within the hydrogenosome, malate and pyruvate were additionally supplied to both monoxenic and axenic cultures alongside the labeled glucose. We anticipated that in monoxenic culture with methanogens, AGF would preferentially follow the malate pathway, leading to increased production of H_2_, CO_2_, and formate. In contrast, axenic was expected to maintain a balance between cytosolic and hydrogenosomic pathways.

## 2. Material and methods Enrichment and Isolation of AGF

Fresh fecal samples of domesticated Zebu cattle (*Bos taurus indicus*) were collected from the Zoo Rapperswil, Switzerland in April 2023. The samples were processed for enrichment and isolation by using the protocol described by Joshi et al. using a modified NRF (no rumen fluid) medium[20]. For isolation, the same medium was used with 5 g L^-1^ cellobiose as a substrate as described before [21].

### 2.1 Morphological and Molecular identification

In a preliminary analysis, by visual inspection and microscopic examination, the morphological traits of the isolate were studied using an Olympus BX53 microscope, equipped with U-FBVW FL filter set (Excitation: 400-440 nm, beam splitter: 455 nm, emission: 460 nm). For molecular identification, the genomic DNA was extracted using the Plant/Fungi DNA Isolation kit (Sigma-Aldrich). The D1/D2 region of Large-Subunit (LSU) rDNA was amplified using NL1/NL4 primers [22] and sequenced at Microsynth (Balgach, Switzerland). GenBank BLASTn was used to perform the sequence similarity search. For the identification of methanogenic partners growing in axenic culture, the same genomic DNA extraction method was used to amplify the archaeal 16S region using the primer pair 86f/1340r, and the same procedure was followed for identification [13]. The D1/D2 region sequence of AGF strain RWM43N2 and the 16S rDNA sequence of its associated methanogen obtained in this study were deposited in NCBI database under the accession numbers OR892617 and OR884089, respectively.

### 2.2 Metabolic Pathway Analysis

The experiment was conducted in a No Rumen Fluid (NRF) medium dispensed in long serum bottles with a total volume of 56.6 mL, sealed with gas-proof butyl rubber stoppers. To assess the preferred metabolic pathway axenic cultures of *Cyllamyces* sp. RWM43N2 and monoxenic cultures with *Methanobrevibacter olleyae* RWM43N3 were fed with different combinations of carbon sources. Three experimental sets were prepared (Figure 1): Set A: Cultures were fed with uniformly ^13^C_6_-labeled glucose at 2 g L^-1^ (11.1 mM).

**Figure 1:**
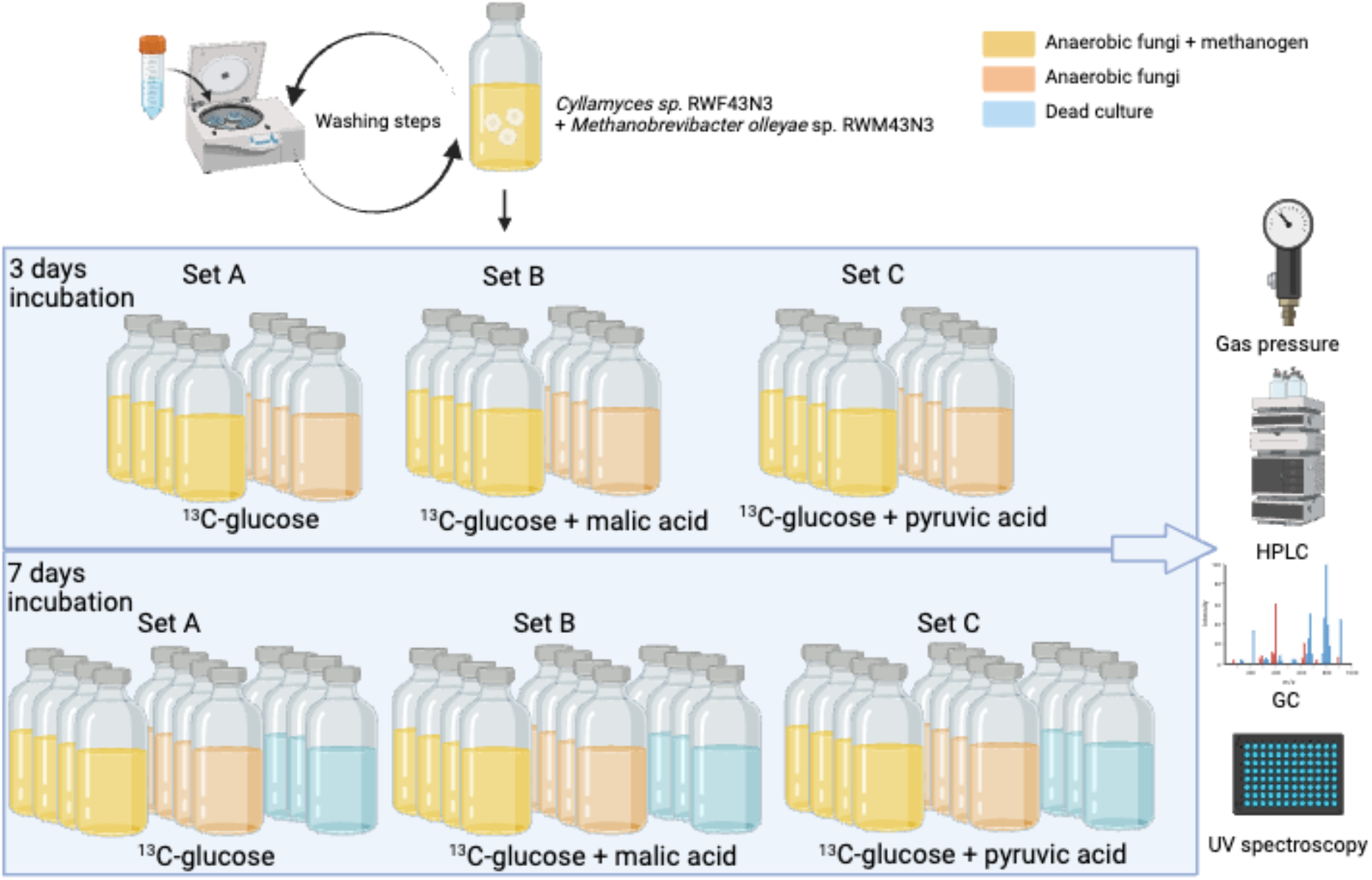
Schematic depiction of the two-stage experimental setup. The top row illustrates the effect of early incubation, where initial bottles were first analyzed after three days of inoculation. The bottles in the bottom row were analyzed seven days after the inoculation. In this stage, a dead control was included to provide a comparative baseline. On the right side, are shown the various methods used to monitor growth and metabolism.

Set B: Cultures contained only ^13^C_6_-labeled glucose at 1 g L^-1^ (5.55 mM) and 15 mM of unlabeled malic acid. The filter sterilized malic acid stock was added just before the inoculation to ensure its stability.

Set C: Cultures contained uniformly ^13^C_6_-labeled glucose at 1 g L^-1^ (5.55 mM) and 15 mM of unlabeled pyruvate. Again, the filter sterilized pyruvate was added just before inoculation. Based on previous reports, the distribution of intermediates between the cytosol and the hydrogenosome was calculated per mole of glucose. To equalize hydrogenosomal substrate supply across all experimental conditions, sets B and C were supplemented with malate and pyruvate. These amounts were calculated based on published cytosol–hydrogenosome intermediate distribution ratios to match the theoretical yield of the omitted 1 g of glucose [9,10,23]. Additional unlabeled malate and pyruvate were supplemented to track the products produced from the metabolism of labeled ^13^C glucose or spiked, unlabeled intermediate metabolites, going to the hydrogenosome.

In order to restrict bacterial growth, the media contained an antibiotic cocktail comprising penicillin G sodium salt (200 μg L^-1^), streptomycin sulfate (200 μg L^-1^), and ampicillin sodium salt (100 μg L^-1^). To ensure sufficient growth in the absence of rumen fluid, the minimal medium contained a 0.1% v/v vitamin solution [20].

Each experimental set comprised four replicates. Prior to inoculation, the biomass was centrifuged at 4000×g for 10 min at RT, and the settled biomass was washed once in a glucose-free medium to remove any unlabeled glucose. To keep the uniformity of AGF inoculum, every bottle was inoculated with 2 mL of washed monoxenic culture. Axenic culture was prepared by adding 0.1% v/v 2-bromoethanesulphonate (BES, 200 μg ml^-1^ final concentration) to suppress the activity of associated methanogen. In negative controls both, BES and cycloheximide (100 μg ml^-1^ final concentration) were incorporated to prevent growth of methanogens and AGF. Additionally sterile medium was added in set A to make the total volume up to 20 ml. One set of four replicates was assessed for growth and end-products after three days of incubation at 39 ^°^C, while the second set of four replicates was kept for prolonged incubation and analyzed after seven days of incubation. This should allow to track potential shifts in metabolism at different growth phases. Control culture bottles, where activity of AGF and methanogen was suppressed using BES and cycloheximide, were only analyzed after seven days. (Figure 1).

### 2.3 Growth and end-product assessment

To accurately measure the volume of produced gas, the total volume, of each bottle was measured prior to the start of the experiment (Table S1). After inoculation, samples were withdrawn (1 mL) from every bottle for 0 hour reading for accurate end-metabolite production. After 3 d and 7 d of incubation, the relative gas pressure was measured using a digital pressure gauge (LEO2, Keller Pressure) equipped with a sterile filter (0.25 μM) and needle at ambient temperature (∼25 °C). Gas chromatography was subsequently conducted to analyze the headspace gas composition. Following gas chromatography, a 2 mL liquid sample was withdrawn after thorough mixing, centrifuged at 11,000 × g and the supernatant was used for HPLC analysis according to [24]. The residual biomass was resuspended in phosphate buffer (100 mM, pH 7) for quantification of AGF and methanogens using UV spectrophotometry as described by Leggieri et al. [25]. The remaining sample was centrifuged at 9,000 × g, supernatant was filtered through a 0.2 μm filter to perform nanoscale high-performance liquid chromatography–high-resolution mass spectrometry (HPLC–HRMS) to quantify labeled end-products, pyruvate, and malate. For this, an ACQUITY UPLC BEH Amide Column (130Å, 1.7 µm, 2.1 mm X 100 mm) served as the stationary phase, operating at 80 ^°^C. The mobile phase was composed of 70 mM ammonium acetate pH 7.4 in 70% acetonitrile solvent, while solvent B contained 70 mM ammonium acetate pH 7.4 in 30% acetonitrile solvent. The eluent followed a multistep gradient: Solvent A 0 min, 0%; 1 min, 50%; 36 min, 90%; 3min, 50%, and for solvent B 0 min, 50%; 1 min, 50%; 3 min, 50%. The sample injection volume was 1 μL and the flow rate was 0.6 μL min^−1^. Average labeling was computed through the selective integration of isotopic peaks and subsequently adjusted for natural labeling as described in Meyer et al. [26]. The residual biomass was subjected to overnight drying at 70^°^C in a bespoke aluminum tray for total biomass quantification.

### 2.4 Statistical analysis

Comparative analysis of gas pressure measurements and UV-spectroscopy data were conducted using Two-way analysis of variance (ANOVA) using GraphPad version 10.0.0 for Windows (GraphPad Software, Boston, MA, USA) in positive and negative control groups across all inhibitors and promoters treatments with positive and negative conditions were evaluated, including the interaction between the two factors. Statistical significance was defined as p<0.05. Prior to ANOVA, the data were assessed for normality using the Shapiro-Wilk test and for homogeneity of variance using Brown-Forsythe test. Where significance differences were detected, Tukey’s multiple-comparison test was used for post-hoc comparison. Data are presented as mean ±SD/SEM from (n) independent biological replicates.

## 3. Results

### 3.1 Morphological and molecular identification

Visual inspection revealed glossy colony morphology and non-uniform, non-filamentous colonies in the roll bottles. The growth in soluble medium displayed a sedimenting, granular, powdery growth, characteristic of bulbous representatives of AGF. The BLAST search of LSU sequences (700–750 bp) of strain RWF43N3 (accession number OR892617) showed 99.6% similarity with *Cyllamyces* sp. strain BFK346. Additionally, the results for archaeal 16S rRNA gene sequences (1200 bp) of strain RWM43N3 (accession number OR884089) revealed 99.89% similarity with *Methanobrevibacter olleyae* strain KM1H5-1P.

### 3.2 Gas pressure measurement

After three days of incubation, the gas production in axenic cultures (C3) surpassed that of monoxenic cultures (CM3) when supplemented with glucose and pyruvate as substrates (Figure 2). Both, axenic and monoxenic cultures, supplied with malate (set B) as a supplementary substrate produced more gas than cultures supplied with pyruvate (set C) or glucose only (set A).

**Figure 2:**
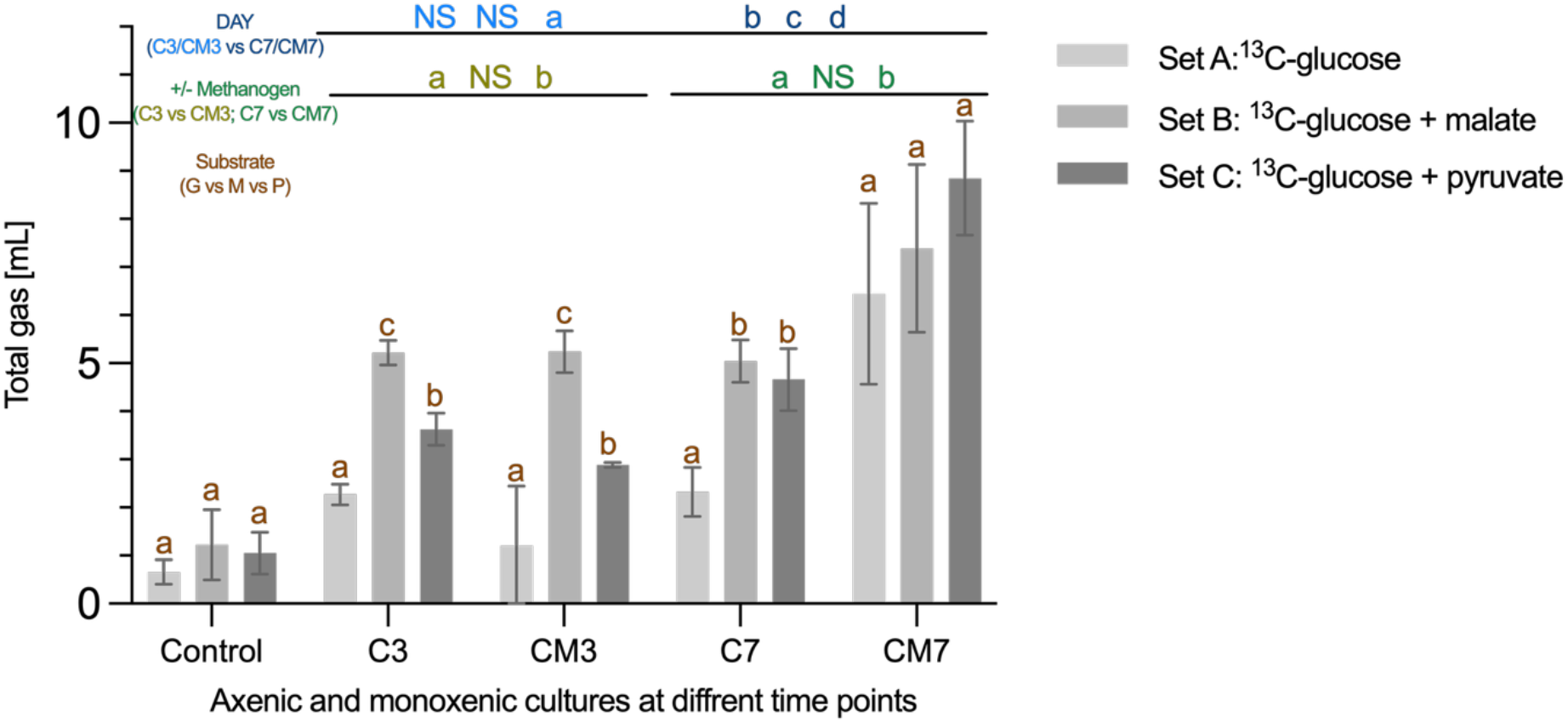
Comparison of gas production (mL) by axenic (**C**) and monoxenic culture (**CM**) of AGF and methanogen grown in different substrates after three days (**C3/CM3**) or seven days (**C7/CM7**). The color-coded bars represent different experimental sets (supplemented with only ^13^C-glucose **(A)**, ^13^C-glucose and malate **(B)**, ^13^C-glucose and pyruvate **(C)** respectively), as described in section 2.2. Error bars indicate standard deviation. Treatment differences (ANOVA) at P<0.05 and P<0.001 are indicated by * and **** respectively. Lowercase letters above bars denote statistically significant differences within each comparison group, color-coded as follows: brown letters compare substrates (glucose vs. malate vs. pyruvate) within the same day and culture type; blue letters compare day 3 vs. day 7 within the same substrate and culture type; green letters compare axenic vs. monoxenic cultures within the same substrate and day. Bars sharing the same letter are not significantly different; “NS” indicates no significant difference.

In contrast, after seven days of incubation, monoxenic cultures produced more gas than in axenic across all substrates. Among axenic cultures, those spiked with malate (B) did not show any difference in gas production after three days of incubation. While all monoxenic cultures growing on different substrates yielded higher gas production compared to their respective axenic cultures. For monoxenic cultures at day 7, highest gas production was observed in treatment supplemented with pyruvate (set C), followed by malate (set B), and with labelled glucose alone (set A) producing the least (Figure 2).

### 3.3 Biomass quantification

Optical density measurement at 450 nm indicated that the growth of AGF was enhanced when it was cultivated in an axenic culture with glucose supplementation compared to a monoxenic culture (Figure 3 A and B). However, supplementation of malate boosted the growth of AGF in monoxenic culture. Contrarily, the addition of pyruvate (C) appeared to enhance the growth of *Cyllamyces* sp. axenic culture but did not show any positive effects on monoxenic culture (Figure 3 A and B).

**Figure 3:**
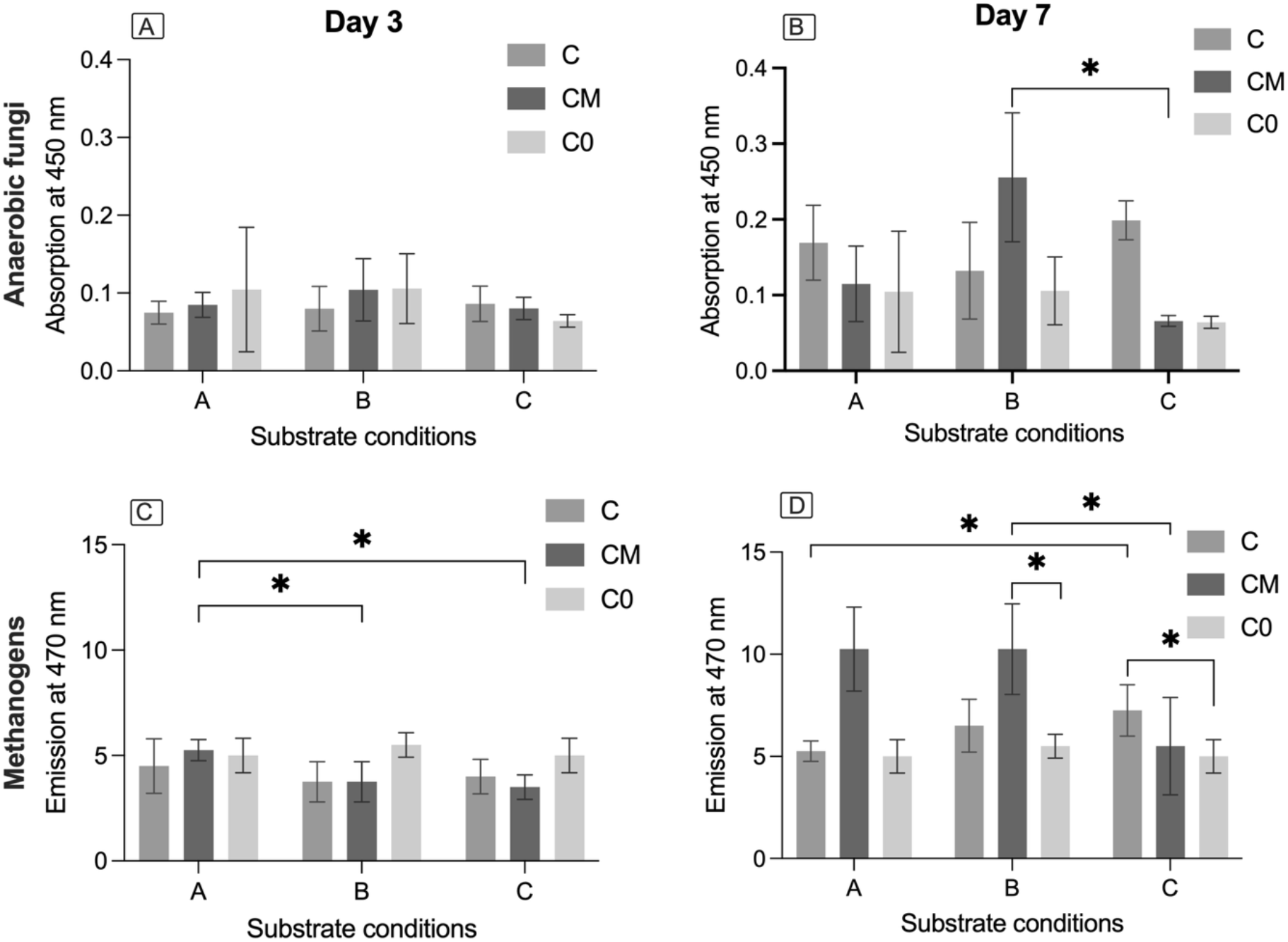
Growth quantification of anaerobic gut fungi and methanogens in both axenic (C) and monoxenic (CM) culture, after incubation of three (left) and seven days (right) of incubation using UV spectrophotometry [3]. The color code distinguishes between axenic culture (C), monoxenic culture (CM) and timepoint 0 values of bottles inoculated with dead culture (C0). The symbols (*) showing the significance level from * to ****, where * = P ≤ 0.05; ** = P ≤ 0.01; *** = P ≤ 0.001; **** = P ≤ 0.0001.

Results of UV spectrophotometry revealed enhanced growth of methanogens when monoxenic cultures were fed with glucose and malate, whereas the addition of pyruvate did not yield a significant impact on the growth of methanogens (Figure 3 C and D).

### 3.4 End-product analyses

Overall monoxenic cultures showed higher glucose utilization after 7 days than axenic cultures, regardless of the carbon sources provided.

#### Set A: growth in ^13^C-glucose

The concentration of ethanol increased significantly after three days of growth, with a more pronounced change observed after 7 days of incubation in monoxenic cultures compared to the axenic cultures (Figure 4A). Gradual formate production was observed in axenic cultures, whereas no formate was accumulated in monoxenic cultures. Low acetate concentrations were observed after three days in both mono-and co-cultures, but after seven days, the concentrations were threefold higher in monoxenic cultures compared to axenic.

**Figure 4:**
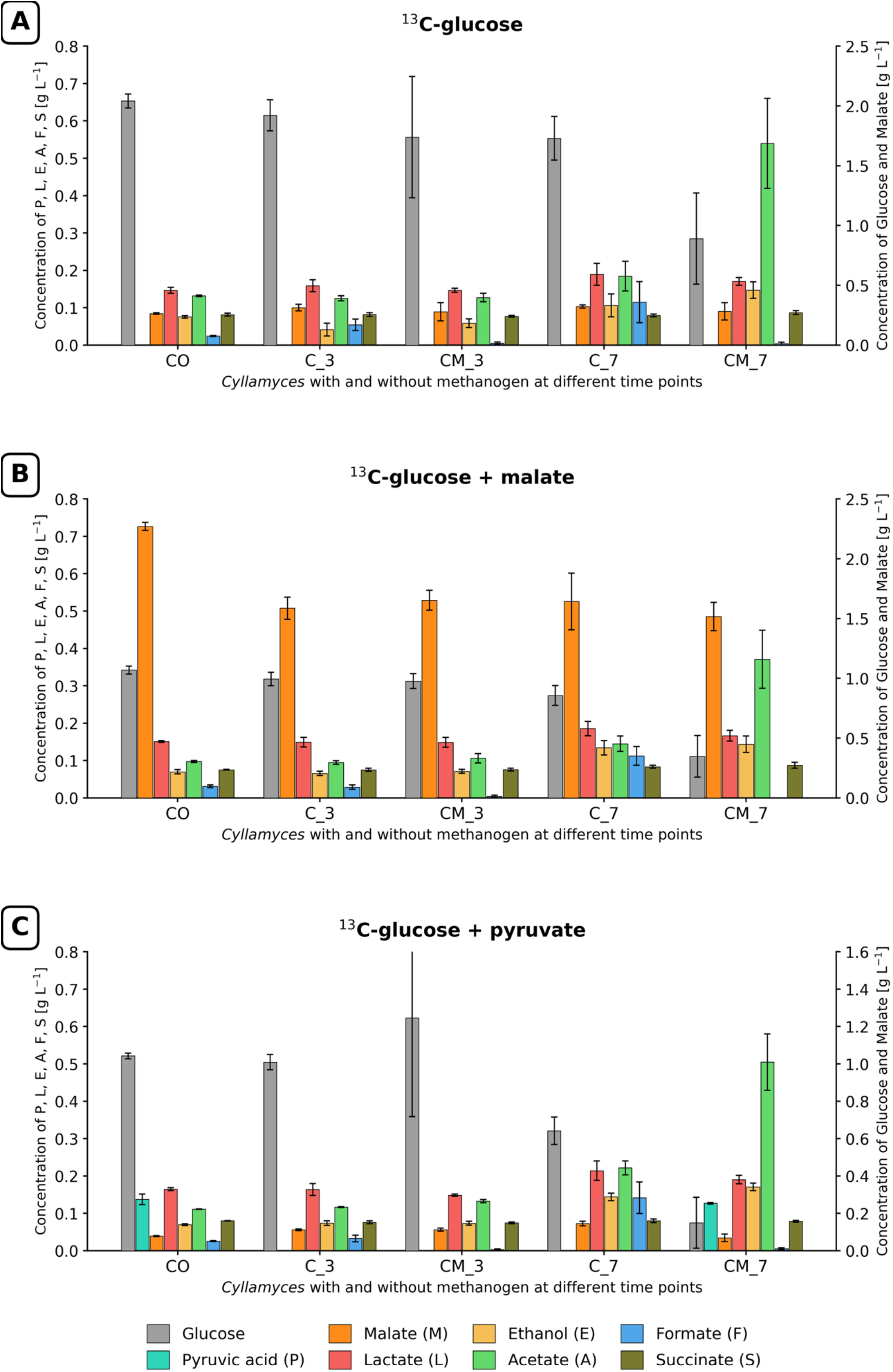
Substrate and metabolite concentrations for axenic (C) and monoxenic (CM) cultures fed with different carbon sources: set A (labeled glucose), set B (labeled glucose + malate), and set C (labeled glucose + pyruvate). Time points three and seven days of incubation are illustrated in numbers. Concentrations of end-products are shown on the left y-axis, while the right y-axis displays ^13^C-glucose in all sets, and malate and pyruvate in sets B and C, respectively.

Additionally, succinate and lactate production did not show any significant difference between the control and treatments over seven days of incubation.

#### Set B: ^13^C-glucose and malate as substrate

Malate concentration was higher in control bottles, equivalent to added malate concentration, and was equal in monoxenic and axenic cultures after three and seven days (Figure 4 B). In axenic culture, there was a gradual increase in formate production after three and seven days of incubation. Ethanol production was higher after seven days of incubation and remained relatively consistent in both monoxenic and axenic cultures after three and seven days of incubation. Lactate production was relatively high in axenic culture after seven days of growth. In contrast, acetate production was 2.5 times higher in monoxenic than in axenic culture.

#### Set C: ^13^C-glucose and pyruvate as substrate

Compared to the glucose (set A) and glucose-malate supplementation (set B), more ^13^C-glucose was utilized in monoxenic culture after seven days of incubation in glucose-pyruvate supplemented set C. Pyruvate concentration was lower in control bottles and monoxenic than in axenic culture after seven days of growth. In fact, it was hardly detectable in three-day-old and seven-day-old axenic cultures (Figure 4C). Unexpectedly, pyruvate concentration in control bottles was lower than the initially supplemented amount (15 mM). Ethanol production was slightly higher in monoxenic than in axenic cultures after seven days of growth. Formate production became noticeable after seven days of growth in axenic culture. Like sets A and B, acetate production in monoxenic culture was nearly double than in axenic culture.

### 3.5 Labeled fraction of pyruvate and malate

Set B, which was spiked with 15 mM unlabeled malate exhibited as expected a very low labeled fraction of malate compared to Set A and C while the overall concentration of malate is elevated in comparison with the other sets. In set A and set C, axenic cultures showed a slightly higher labeled malate fraction compared to monoxenic cultures. Moreover, the labeling fraction of malate increased from 3 days to 7 days, while almost no labeled malate fraction was seen in the dead culture control bottles throughout the whole experiment. In all the sets the concentration of malate is slightly lower in the case of monoxenic compared to axenic cultures and more pronounced at 7 days (Figure 5A).

**Figure 5:**
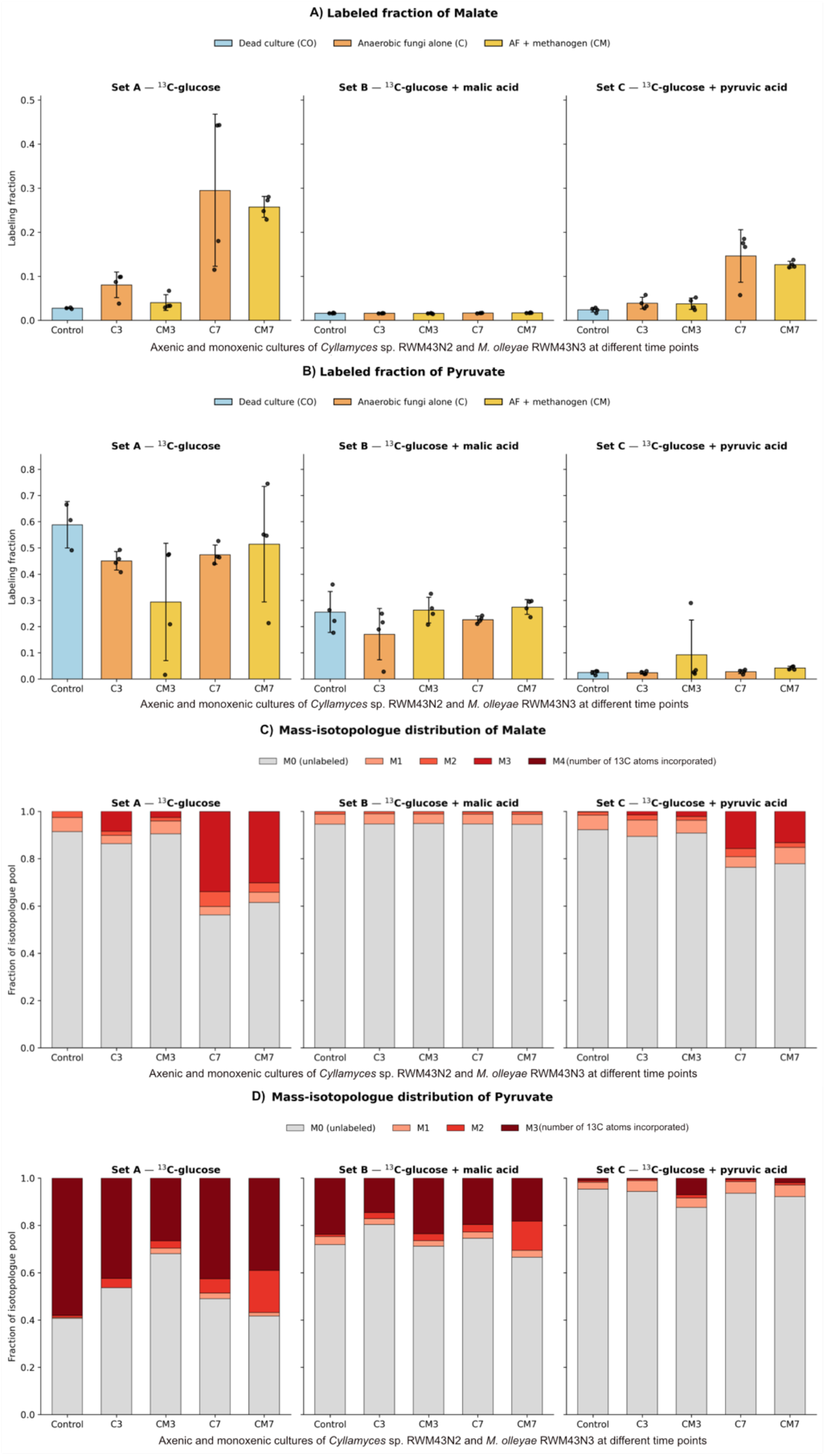
^13^C labelling of malate and pyruvate in axenic and monoxenic cultures of *Cyllamyces* sp. RWM43N2 and *M. olleyae* RWM43N3. Shown are the labelled fractions of malate (A) and pyruvate (B), and the mass-isotopologue distributions of malate (C) and pyruvate (D), across three experimental sets: Set A, ^13^C-glucose; Set B, ^13^C-glucose + malic acid; and Set C, ^13^C-glucose + pyruvic acid. Samples were taken from axenic fungal cultures (C) and monoxenic fungus–methanogen co-cultures (CM) after 3 and 7 days (C3, CM3, C7, CM7). A time-0 control inoculated with dead culture (CO) is also shown. In A and B, bars show the mean ± standard deviation of four replicates, with individual replicates shown as points. The y-axis scales differ between A and B for better visualization. In C and D, stacked bars show the mean fractional abundance of each isotopologue, where M0 is unlabelled and Mn indicates the number of ^13^C atoms incorporated (up to M4 for malate and M3 for pyruvate).

Similarly, pyruvate spiked set C showed almost no labeled pyruvate, while in sets A and B the pyruvate labeling remained relatively constant for all samples throughout the experiment, higher in set A compared to set B. The overall concentration of pyruvate is elevated in set C compared to sets A and B, while in all the sets pyruvate concentrations tend to be higher in the case of the monoxenic compared to the axenic cultures (Figure 5B).

## 4. Discussion

The molecular identification revealed the syntrophic association of *Cyllamyces* sp. with *Methanobrevibacter olleyae* which confirmed the coherent syntrophic association of AGF with *Methanobrevibacter* [27]. This study demonstrated the influence of associated methanogens on the activity and growth of AGF, via monitoring of gas production and biomass formation (Fig. 2 and 3). The syntrophic association with methanogens is proven to influence AGF activity, specifically gas production, enzyme activity, and substrate utilization [8,10,27,28]. This is due to the shifting electron sink from the cytosolic towards the hydrogenosomic pathway as mentioned in Li et al. [26]. This study also noted the difference in substrate utilization pattern of AGF in the presence of methanogens. In the initial three days of growth, malate supplementation was observed to promote the growth of both, the axenic and monoxenic cultures, as confirmed by higher gas production than with glucose supplementation alone. Monitoring of total gas measurement is considered to be a non-destructive and reliable method for AGF growth evaluation [29]. This also refers to the sign of active syntrophy with methanogens. However, UV spectrophotometry did not support this observation, possibly due to the detection limit caused by inadequate growth of AGF and methanogens, which may not have differed significantly from the control. This may have been caused by the detection limit, as well as interference from dead biomass in control bottles.

Contrarily, after seven days of growth, the growth of AGF and methanogens in malate-supplemented monoxenic culture exhibited a mutual symbiosis in terms of biomass formation as depicted in Figure 2b., which partially supports the hypothesis of this study that the malate pathway aids methanogens’ growth through H_2_ and CO_2_ production. Additionally, AGF also formed more biomass, aligning with the findings of Marvin-Sikkema et al. [30] on *Neocallimastix* sp. L2 cultures and showcasing the vital role of malate in energy metabolism. This implies that the generation of H_2_ via reduced ferredoxin, as previously outlined [30], occurs through the activity of hydrogenase and PFO enzymes, facilitating H_2_ production from reduced ferredoxin even within environments with high H_2_ concentrations. Furthermore, this process contributes to the preservation of cellular redox potential by utilizing NADPH through reduction mechanisms.

Unexpectedly, monoxenic culture supplemented with pyruvate exhibited the highest gas production but also displayed low biomass formation of both *Cyllamyces* sp. and *M. olleyae*. This suggests that higher CO_2_ to H_2_ was less favorable for methanogenesis via spiked pyruvate pathway (Figure 6). This leads to controversial pyruvate to acetyl Co-A conversion via PFL (pyruvate formate lyase) or PFO (pyruvate-ferredoxin oxidoreductase) leading to *the* formation of either formate or CO_2_, respectively (Figure 6). Unlike in Boxma et al. [19], the axenic culture of *Cyllamyces* sp. supplemented with glucose only did not show 1:1:1 formate, ethanol, and acetate production, rather it produced 1:1 ethanol and formate, while double acetate, suggesting involvement of PFO in pyruvate to acetyl Co-A conversion, which is less active in *Neocallimastigomycota* [9,19,30]. This higher gas production in monoxenic culture spiked with pyruvate leads to more H_2_ and CO_2_ formation via PFO or PDH (pyruvate dehydrogenase) seems to be thermodynamically less favorable for methanogenesis. Here, less biomass formation of *Cyllamyces* sp. could be explained by less recycling of NADP, which occurs via malate metabolism in the hydrogenosome [30]. This was again corroborated by high acetate formation in monoxenic cultures spiked with pyruvate. This also leads to another discussion on the presence of either PFO or PFL in different species of AGF, as suggested by Wilken et al. [9].

**Figure 6:**
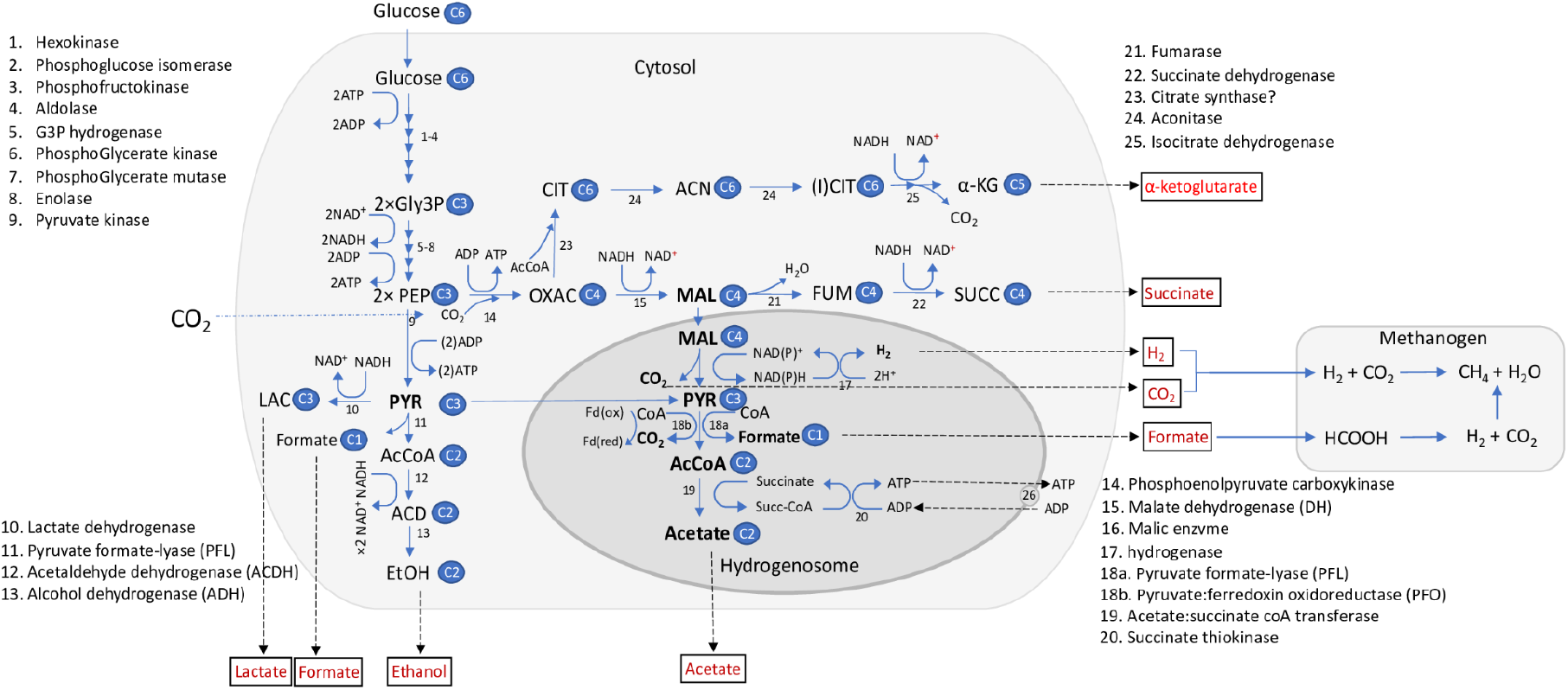
Proposed model of the biochemical pathway of anaerobic gut fungi in both the cytosol and the hydrogenosome, with glucose as initial substrate. The enzymes involved in the reactions are depicted with the respective numbers and corresponding annotations within the biochemical pathways. The short form in the diagram refers to Gly3P, glyceraldehyde 3-phosphate; PEP, phosphoenol pyruvate; PYR, pyruvate; LAC, lactic acid; AcCoA, acetyl co-enzyme A; ACD, acetaldehyde; EtOH; ethanol; OXAC, oxalic acid; MAL, malic acid, FUM, fumarate; SUCC, succinate; CIT; citric acid; ACN, cis-aconitic acid, (I)CIT, iso-citric acid, α-KG, α-ketoglutaric acid.

Axenic culture supplemented with pyruvate, exhibiting more gas production compared to a glucose-fed axenic culture, this might be attributed to H_2_ and CO_2_ generation via PFO. Moreover, the observed imbalance in acetate, ethanol, and formate production (Figure 4c) indicates that surplus acetate might have been coming from supplemented pyruvate. This reconfirms the involvement of the PFO in this process. The malate-supplemented axenic culture exhibited more gas production (Figure 2) including CO_2_ and H_2_ via malic enzyme and hydrogenase, while less biomass formation (Figure 3b).

The glucose metabolism appeared to promote methanogen growth (Figure 3D) by initiating the natural syntrophic cycle where the metabolism is shifted towards the hydrogenosomic pathways. This primarily occurs through redirecting the electron flow towards the malate pathway, as evidenced in this study, under malate supplementation. Surprisingly, this shift resulted in reduced biomass formation in AGF. In contrast, *Cyllamyces* sp. axenic culture showed increased biomass formation compared to monoxenic culture. This implies that monoxenic cultures channels more electrons flux towards the hydrogenosome, likely via the malate pathway, thereby increasing the supply of H_2_ for methanogenesis. This is in agreement with reduced malate concentrations in the monoxenic culture compared to the axenic. Here again, pyruvate concentration showed the opposite trend (Figure 4C). The syntrophic interaction with methanogens redirects electron sink in AGF towards the hydrogenosome [10], increasing energy availability for substrate breakdown and aiding in the provisioning of substrate for methanogenesis. As a result, this mechanism reduces fungal biomass production. While this likely regulates fungal biomass growth in their natural habitat, its effects in practical applications require detailed investigation.

Both pyruvate:ferredoxin oxidoreductase (PFO) and pyruvate-formate lyase (PFL) are crucial enzymes involved in the conversion of pyruvate in AGF [31]. The specific involvement of PFO or PFL in *Neocallimastigomycota*-hydrogenosomes has been debated. PFO contributes to the conversion of pyruvate to acetyl Co-A and CO_2_, utilizing ferredoxin as an electron carrier in H_2_ formation. While PFL generates formate as a byproduct. Several studies supporting the involvement of PFL in the hydrogenosome based on the activity measurement; however, these findings lack concrete evidence and remain contentious, since the possibility of cytosolic PFL activity cannot be entirely excluded [32,33]. In contrast, relatively few studies have supported the involvement of PFO in hydrogenosomic conversion of pyruvate, with recent research suggesting that either enzyme’s involvements may vary across different genera of AF. The involvement of PFO or PFL in the hydrogenosomic metabolism may influence mutual benefits for syntrophic methanogens, as also corroborated in this study. PFL-dependent metabolism in hydrogenosomes results in the production of H_2_ through malic enzyme and hydrogenase, along with formate and CO_2_ produced by PFL. In contrast, PFO-dependent metabolism primarily produces H_2_ and CO_2_ to be used by methanogens [30]. Although many methanogens have high affinity for H_2_, due to thermodynamic benefits, formate can serve as an alternative interspecies electron carrier preferred by methanogens within syntrophic environments and can be readily utilized by many methanogens [31].

Future studies exploring the syntrophic association should delve into the impact of hydrogenosome metabolism via PFL or PFO, evaluating the influence on H_2_ and CO_2_ production and their indirect effects on methanogenesis.

## 5. Conclusion

This investigation sheds light on the intricate syntrophic association between *Cyllamyces* sp. and *Methanobrevibacter olleyae*, emphasizing the substantial impact of associated methanogens on AGF. The findings revealed the inclination of *Cyllamyces* sp. towards the malate pathway, supporting the growth of an associated methanogen, *Methanobrevibacter olleyae*. In association with M. *olleyae, Cyllamyces* sp. exhibited enhanced growth and favored the malate pathway in the hydrogenosome. Furthermore, the study suggests the involvement of pyruvate:ferredoxin oxidoreductase (PFO) instead of pyruvate formate lyase (PFL) in pyruvate metabolism, that results in increased gas production. Pyruvate supplementation exhibited enhanced fungal biomass formation, indicating its favorability in AGF axenic culture. However, the syntrophic association with *M. olleyae* led to high activity and less biomass formation in *Cyllamyces* sp. prompting further inquiry into mutual benefits of syntrophic associations between anaerobic gut fungi and their commonly associated methanogenic partners.

## Supplementary Materials

Table S1: Raw data of gas pressure measurement, HPLC, and GC analysis; Table S2: Raw data of labeled fraction analysis; Supplementary figure 1: Differential Interference Contrast (DIC) and fluorescence microscopic images of monoxenic culture.

## Funding

This research was part of a HiPoAF project entitled ‘Unleashing the hidden potential of anaerobic fungi’ and was funded in whole by the Swiss National Science Foundation (SNSF) grant no. 310030E_179552 in Switzerland and the Austrian Science Fund (FWF) [grant 10.55776/I3808].

## Acknowledgments

We extend our gratitude to Roger Fehér for his HPLC analysis and expertise, and Prof. Dr. Mark Van der Giezen (Department of Chemistry, Bioscience and Environmental Engineering, University of Stavanger, Norway) for the insightful discussion on anaerobic metabolism.

## Conflicts of Interest

The authors have no conflicts of interest to declare and there is no financial interest to report.

## Notes

### Competing Interest Statement

The authors have declared no competing interest.

