## Supplementary Figure 1 for "Deciphering efficient metabolic pathways in an anaerobic gut fungus coupled with a syntrophic methanogen"

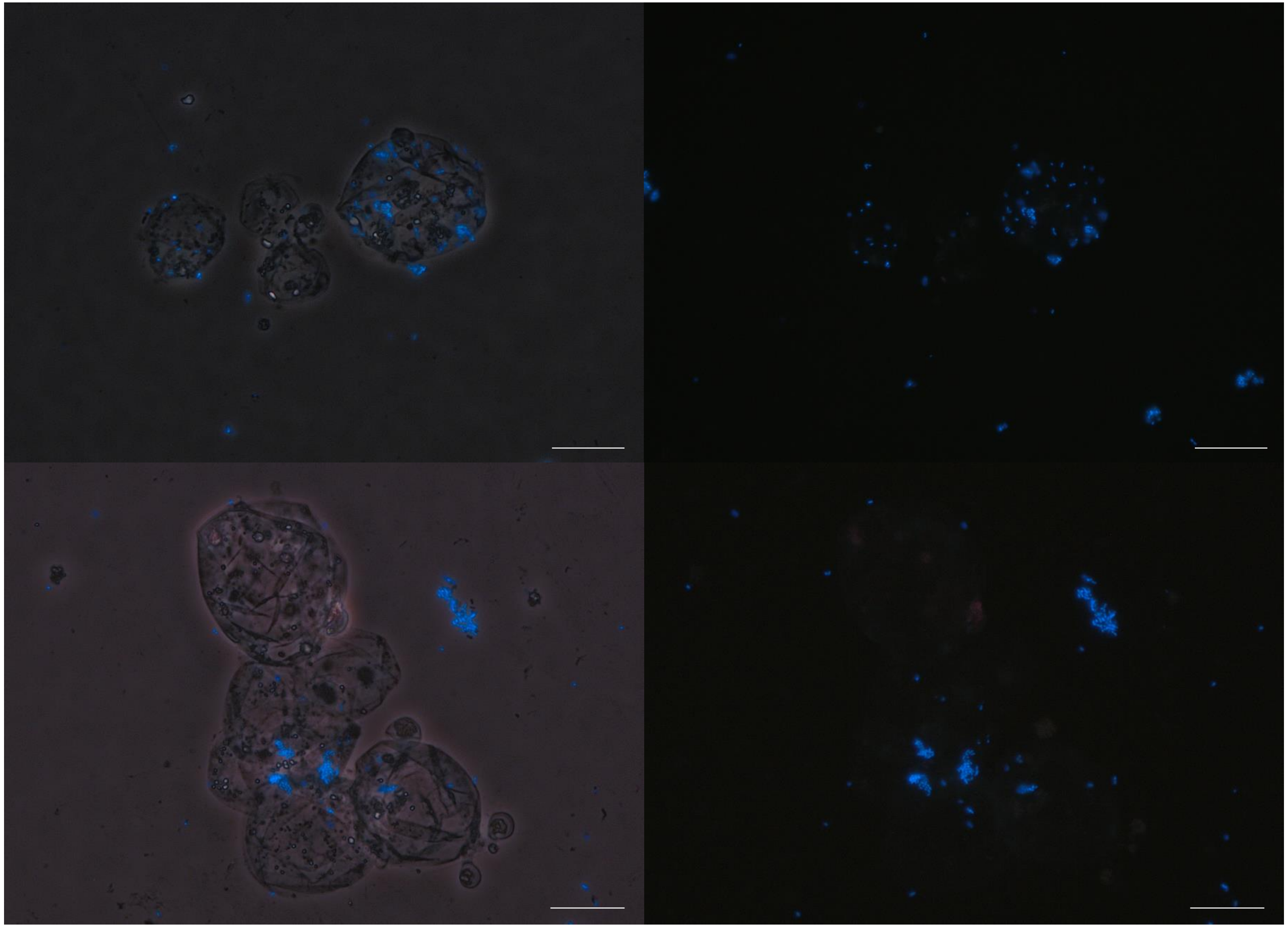

Supplementary figure 1: Monoxenic clture of *Cyllamyces* sp. RWF43N3 and *M. olleyae* RWM43N3 using both Differential Interference Contrast (DIC) microscopy with overlay of UV fluorescence on the left, and only UV fluorescence microscopy on the right. Methanogens (blue) are visible, both adhering to the AGF and in free suspension The scale bar corresponds to 20 µm.
